# Mouse model of SARS-CoV-2 reveals inflammatory role of type I interferon signaling

**DOI:** 10.1101/2020.05.27.118893

**Authors:** Benjamin Israelow, Eric Song, Tianyang Mao, Peiwen Lu, Amit Meir, Feimei Liu, Mia Madel Alfajaro, Jin Wei, Huiping Dong, Robert J Homer, Aaron Ring, Craig B Wilen, Akiko Iwasaki

## Abstract

Severe Acute Respiratory Syndrome-Coronavirus 2 (SARS-Cov-2) has caused over 5,000,000 cases of Coronavirus disease (COVID-19) with significant fatality rate.^1–3^ Due to the urgency of this global pandemic, numerous therapeutic and vaccine trials have begun without customary safety and efficacy studies.^4^ Laboratory mice have been the stalwart of these types of studies; however, they do not support infection by SARS-CoV-2 due to the inability of its spike (S) protein to engage the mouse ortholog of its human entry receptor angiotensin-converting enzyme 2 (hACE2). While hACE2 transgenic mice support infection and pathogenesis,^5^ these mice are currently limited in availability and are restricted to a single genetic background. Here we report the development of a mouse model of SARS-CoV-2 based on adeno associated virus (AAV)-mediated expression of hACE2. These mice support viral replication and antibody production and exhibit pathologic findings found in COVID-19 patients as well as non-human primate models. Moreover, we show that type I interferons are unable to control SARS-CoV2 replication and drive pathologic responses. Thus, the hACE2-AAV mouse model enables rapid deployment for in-depth analysis following robust SARS-CoV-2 infection with authentic patient-derived virus in mice of diverse genetic backgrounds. This represents a much-needed platform for rapidly testing prophylactic and therapeutic strategies to combat COVID-19.

## Development of SARS-CoV-2 mouse model

To overcome the limitation that mouse ACE2 does not support SARS-CoV-2 cellular entry and infection^6,7^, we developed a mouse model of SARS-CoV-2 infection and pathogenesis by delivering human ACE2 (hACE2) into the respiratory tract of C57BL/6J (B6J) mice via adeno-associated virus (AAV9) (Fig.1a). Control (AAV-GFP or mock) and AAV-hACE2 mice were intranasally infected with 1×10^6^ PFU SARS-CoV-2 (passage 2 of isolate USA-WA1/2020). Mice were sacrificed at 2, 4, 7, and 14 days post infection (DPI). During the 14-day time course, mice were monitored daily for weight loss. None developed significant weight changes or died. Compared to control, AAV-hACE2 mice supported productive infection indicated by >200-fold increase in SARS-CoV-2 RNA (Fig.1b) as well as the presence of infectious virus as indicated by plaque assay (Fig.1c).

**Figure 1.**
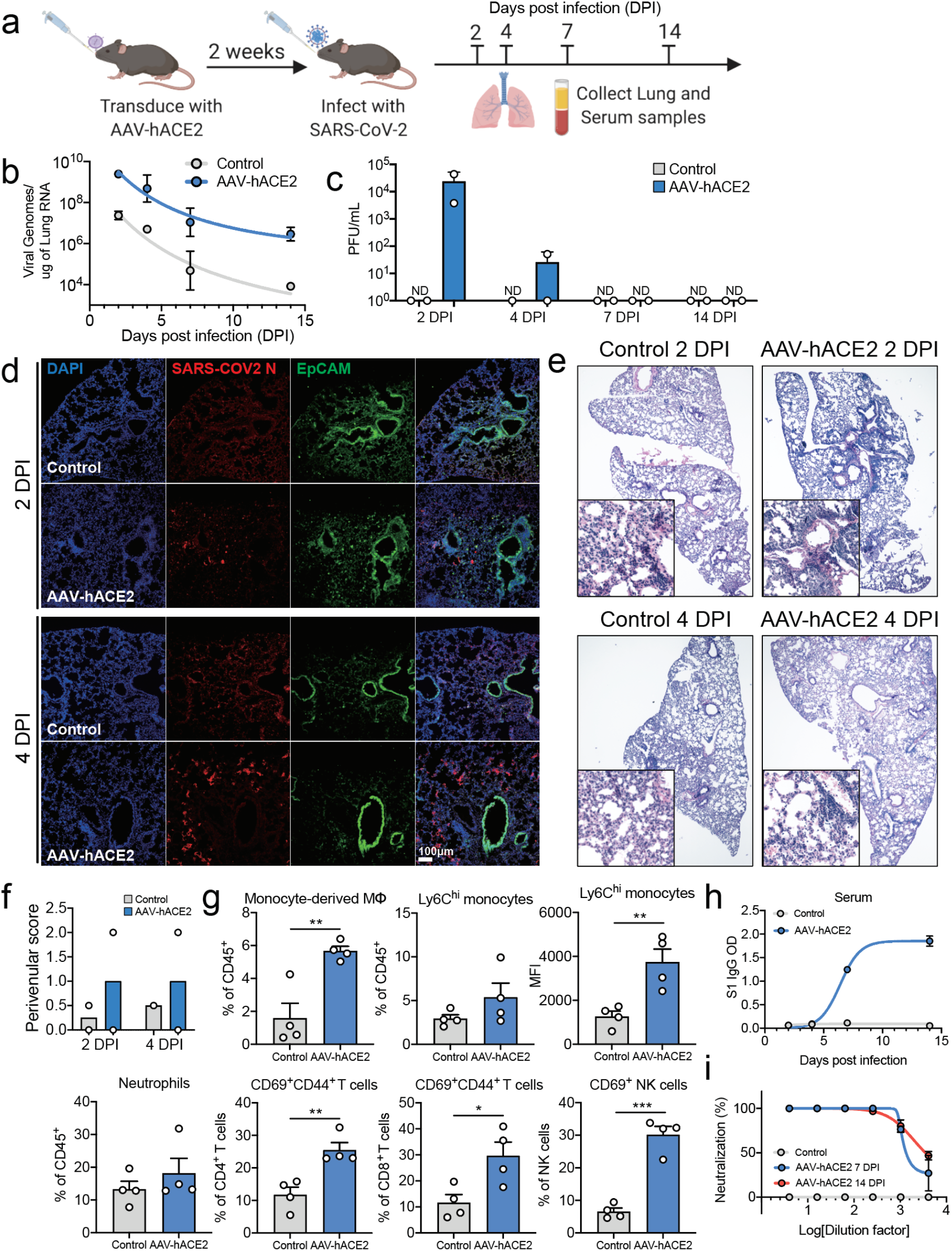
AAV-hACE2 transduction allows for productive SARS-CoV-2 infection *in vivo*. **a,** Schematic of experimental plans. C57BL/6J mice were transduced intratracheally with an adeno-associated vector coding for hACE2 (AAV-hACE2) or control (AAV-GFP or PBS) and infected with SARS-CoV-2 two weeks after. Lung and blood samples were collected at days 2, 4, 7, and 14 days for analysis. **b,** Viral RNA from lung homogenates were measured using qPCR against SARS-CoV-2 N (CDC N1 primers). **c,** Viral titer from lung homogenates were performed by plaque assay on VeroE6 cells. **d,** Frozen lung tissue was stained for SARS-CoV-2 N protein (red) and epithelial cells (EpCAM, green). **e,** Fixed lung tissue was paraffin embedded and stained with H&E. **f**, Images from **e** were scored by a pulmonary pathologist for perivenular score. **g,** At two days post infection, single cell suspensions of lung were analyzed by flow cytometry. Data are shown as frequency of CD45+ cells (monocyte-derived macrophages, Ly6Chi monocytes, and neutrophils), frequency of parent cells (CD44+CD69+ CD4+ T cells, CD44+CD69+ CD8+ T cells, and CD69+ NK cells), or mean fluorescence intensity of CD64 (Ly6Chi monocytes). **h,** Serum antibodies were measured against spike protein using an ELISA. **i,** Day 7 and 14 sera from **h** was used to perform a plaque reduction neutralization assay on VeroE6 cells incubated with SARS-CoV-2.

We next performed histopathologic examination of lung sections from 2- and 4-days post infection (DPI). We found mild diffuse peribronchial infiltrates in AAV-hACE2 mice, which was minimal in control mice (Fig.1e,f). Immunofluorescence staining (Fig.1d) of lung sections revealed diffuse infection (SARS-CoV-2 N protein/Red) within alveolar epithelia (EpCAM/Green). Similar to findings in COVID-19 patients^8^, we found an expansion of pulmonary infiltrating myeloid derived inflammatory cells characterized by Ly6C^hi^ monocytes and inflammatory monocyte-derived macrophages (CD64^+^CD11c-CD11b^+^Ly6C^+^) (Fig 1g; Extended Data Fig. 1d,e). Additionally, we observed relative increases of activated lymphoid cells in lung tissue, including increased percentages of CD69^+^(recent activation) and CD44^+^(recent antigen exposure) CD4^+^ and CD8^+^ T cells (Fig 1g; Extended Data Fig. 1b,c). Lastly, the population of activated (CD69^+^) NK cells also expanded during early infection.

The role of adaptive immunity and specifically antibody response to SARS-CoV-2 is particularly important in the development of safe and effective vaccines. To assess the capacity for B6J AAV-hACE2 mice to mount an antibody response to SARS-CoV-2 challenge, we quantified anti-spike protein IgG titers by ELISA^9,10^. We found that while control infected mice did not develop anti-spike antibodies, AAV-hACE2 B6/J mice mounted a significant antibody response between 4- and 7-DPI, which continued to increase at 14 DPI (Fig. 1h). Next, to assess the neutralization potential of these antibodies, we performed plaque reduction neutralization assay (PRNT) using SARS-CoV-2, and found PRNT75 at a serum dilution of 1:1024 as early as 7 DPI (Fig 1i).

## Interferon stimulated genes and inflammatory cytokines are acutely upregulated during SARS-CoV-2 infection

In a recent study, Blanco-Melo et. al. showed cytokine signatures that are out of proportion to the interferon response in autopsy samples from COVID-19 patients, infected ferrets, and SARS-CoV-2 infected cells in culture^11^. However, others have reported elevated interferon signatures in the lungs of COVID-19 patients^12^. To assess both the cytokine and interferon response to SARS-CoV-2 infected AAV-hACE2 mice, we performed RNA sequencing from infected lung at 2 DPI. In contrast to the control infected mice, AAV-hACE2 mice developed clear signatures of cytokines and interferon stimulated genes (ISGs). In fact, the majority of the differentially expressed upregulated genes were either ISGs or cytokines (Fig. 2a). Interestingly, neither type I, nor type II, nor type III interferons seemed to be upregulated. Using the Interferome database (http://www.interferome.org/), we mapped the top upregulated genes to their transcriptional regulation by either type I, II, or III IFNs (Fig. 2b). We found that while most of the genes shared regulation by type I and type II (and some by all 3 IFNs), a distinct subset of 45 genes that were specific to type I interferon signaling was elevated in the infected lung. We next performed gene ontology analysis of the top upregulated genes, which revealed enrichment of gene clusters in virus-host interaction, immune response, as well as immune cell recruitment and activation (Fig. 2c), consistent with the proinflammatory immune cell infiltrates seen in patients. To analyze the concordance between our model and the recently published patient lung autopsy gene expression from Blanco-Melo et.al., we found that 73% of our shared upregulated genes were ISGs (Fig 2d, e). These results indicated that our mouse model largely recapitulated the transcriptome changes observed in the lungs of COVID-19 patients.

**Figure 2.**
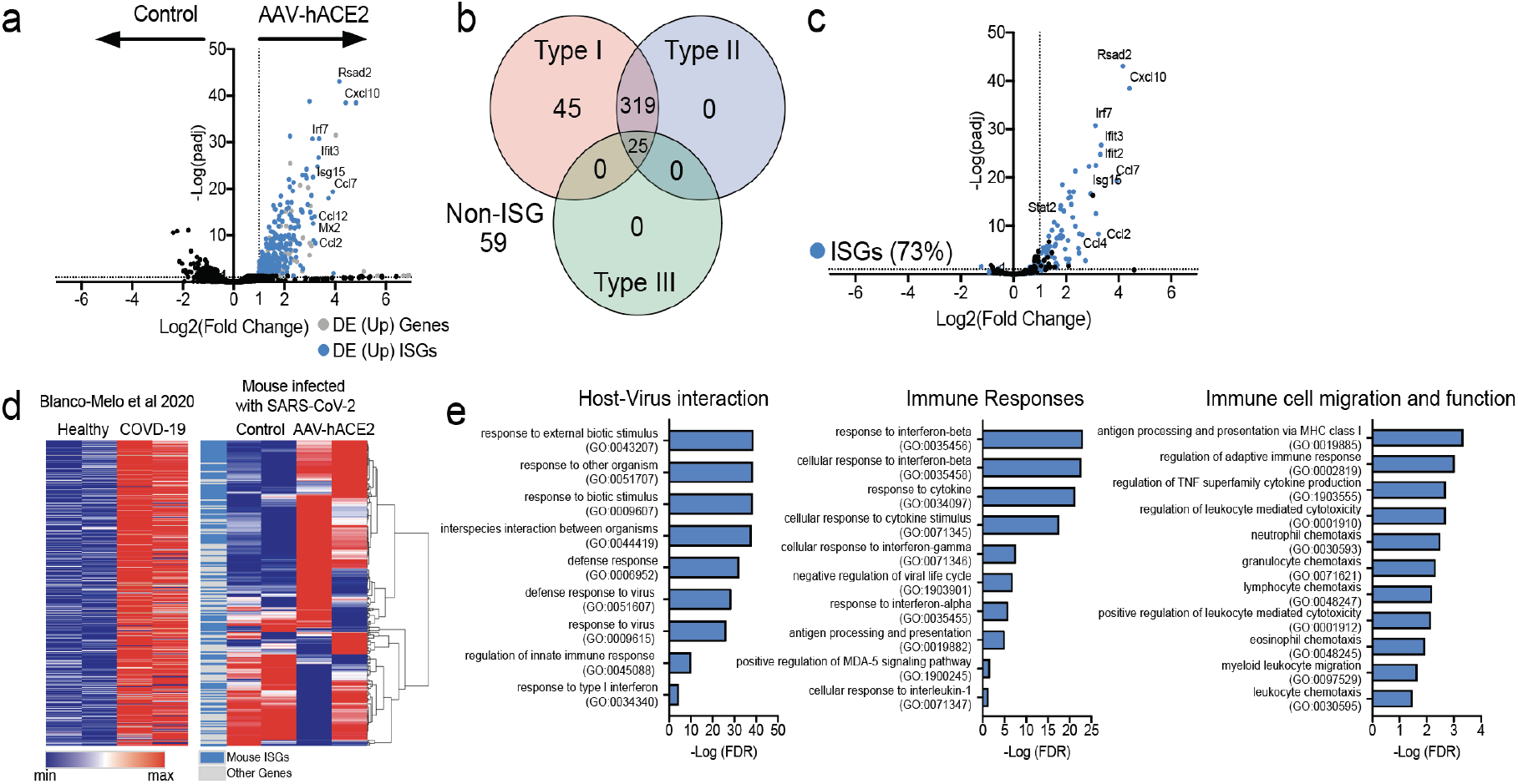
AAV-hACE2 mice infected with SARS-CoV-2 show similar interferon signatures as COVID19 patients. **a,** Volcano plot showing differential expression of genes from whole lungs of mice infected with SARS-CoV-2 with and without AAV-hACE2 at day two post infection. Gray indicates significantly differentially upregulated genes, and blue indicates subsets of genes that are known ISGs. **b,** Significantly upregulated genes were put into Interferome (www.interferome.org) to identify how many genes are stimulated by type I, type II, or type III interferons. Subset of differentially expressed genes in **a** that were significantly upregulated in lungs of COVID19 patients (from Blanco-Melo et al 2020).**, d,** Upregulated gene list from human samples (Blanco-melo et. al. 2020) was graphed (left panel) and used to perform hierarchical clustering for differentially expressed genes from lungs of AAV-hACE2 SARS-CoV-2 infected mice (blue, mouse ISGs; gray, other genes)., **e,** Go Enrichment Analysis was performed on significantly upregulated genes to identify enriched cellular processes.

## Type I interferon signaling is required for recruitment of proinflammatory cells into the lungs and ISG expression, but not for viral clearance

To further investigate the role of type I interferon signaling in SARS-CoV-2 infection, we transduced interferon alpha receptor deficient B6/J mice (IFNAR^−/−^) and interferon regulatory transcription factor 3/7 double knockout B6/J mice (IRF3/7^−/−^) two key transcription factors needed to induce IFNs and ISGs^13^, with AAV-hACE2 and infected them intranasally with 1×10^6^ PFU SARS-CoV-2. We found modest elevations ~10 fold) in viral RNA of IFNAR^−/−^ but not IRF3/7^−/−^ relative to WT B6/J AAV-hACE2 at 2 DPI, though these differences were lost at later time points during infection (Fig. 3a). Viral replication was only slightly enhanced in IFNAR^−/−^ or IRF3/7^−/−^ mice over wild type mice, similar to what has been previously been shown in SARS-CoV-1^14^. In contrast, we found a loss of recruitment of Ly6C^hi^ monocytes and monocyte-derived macrophages in mice deficient in IFNAR or IRF3/7 (Fig. 3d,e). Additionally, we found reduced activation of CD4^+^, CD8^+^, or NK cells in IRF3/7^−/−^ infected mice, and a complete loss of activation of these cell populations in IFNAR^−/−^ infected mice (Fig. 3f-h). In contrast, we observed robust recruitment of neutrophils in the infected IFNAR^−/−^ mice (Fig. 3i). This may reflect the loss of IFN-mediated regulation of inflammasomes leading to neutrophil recruitment^15–17^. These results were consistent with previous reports of pulmonary type I interferon signaling representing a major driver of both recruitment and activation of proinflammatory immune cells in viral infections including SARS-CoV-1^14^ and other respiratory viral infections^18^. RNA sequencing at 2 DPI from the lungs of infected WT, IFNAR^−/−^, and IRF3/7^−/−^ mice reveal markedly elevated ISGs and cytokines profile. Heat map analysis of the top 100 upregulated genes showed significant elevation of antiviral ISGs (Rsad2, IRF7, OASL, Ifit, STAT2, MX2) as well as monocyte recruiting chemokines (Cxcl10, Ccl7, and Ccl2), which were not upregulated in either IFNAR^−/−^ or IRF3/7^−/−^ infected mice (Fig. 3c).

**Figure 3.**
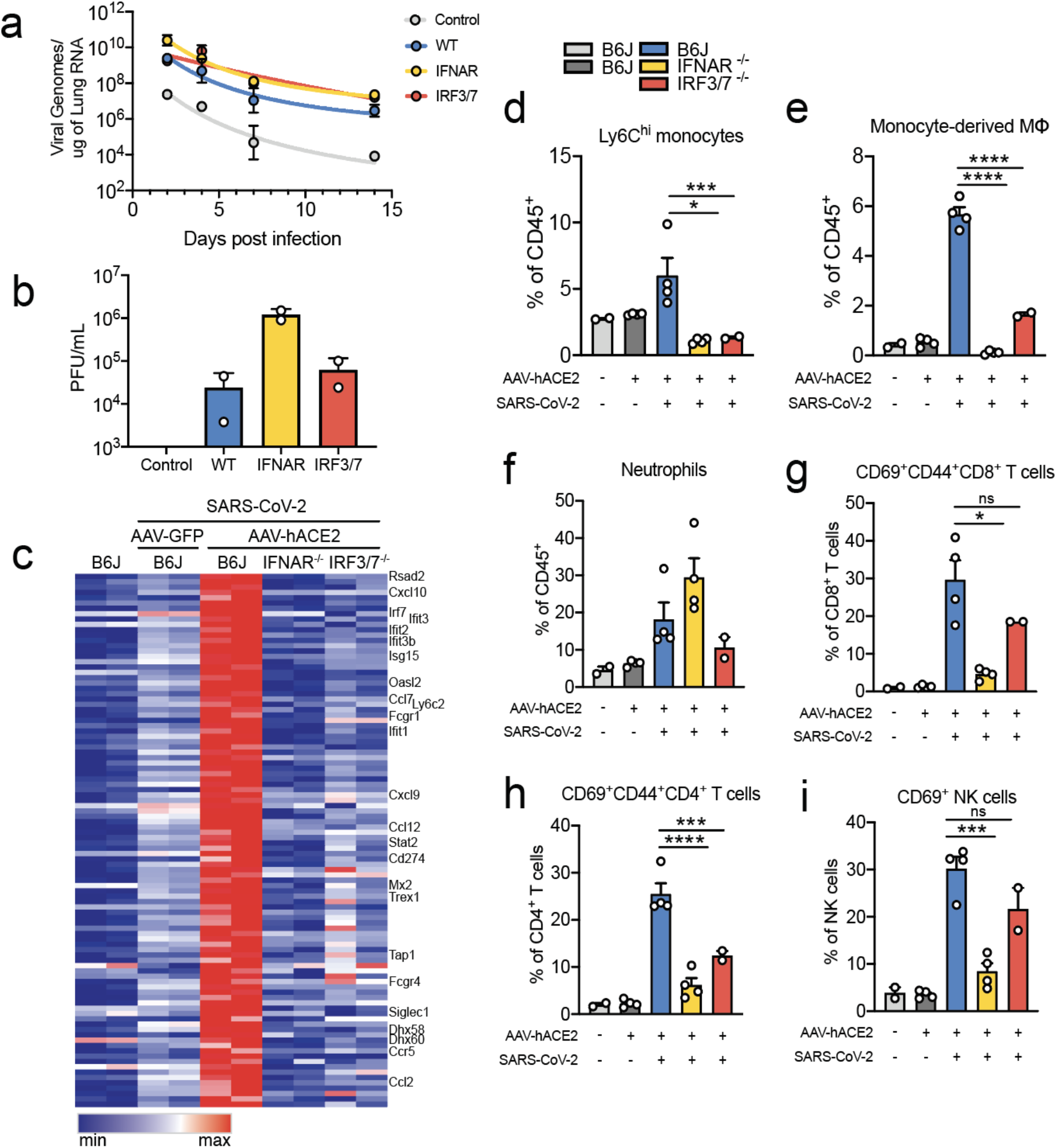
AAV-hACE2 mice infected with SARS-CoV-2 exhibit Type I Interferon dependent immune cell infiltration. C57BL/6J (WT), IFNAR knockout and IRF3/7 double knockout mice were transduced intratracheally with an adeno-associated vector coding for hACE2 (AAV-hACE2) and infected with SARS-CoV-2 two weeks later. **a,** Viral titers in the lung of mice were measured using qPCR against SARS-CoV-2 N (Control and WT were from the same experiment as Figure 1b). **b,** Lung homogenates tittered on VeroE6 cells. **c,** Heat map of top 100 upregulated genes in SARS-CoV-2 infected C57BL/6J mice transduced with AAV-hACE2 versus SARS-CoV-2 infected AAV-GFP transduced mice. **d-i** At day two post infection, lungs of mice were made into single cell suspensions for flow cytometry.

## Discussion

In this study, we describe the development of a mouse model of SARS-CoV-2 using AAV-hACE2 delivery to the mouse respiratory tract. This model develops productive SARS-CoV-2 infection as well as inflammatory pulmonary infiltrates that are characteristic of COVID-19 in humans. We found an acute inflammatory immune response characterized by infiltrating monocytes, macrophages and neutrophils, as well as activated T and NK cells. We also found gene signatures of acute ISG response which clusters most closely with type I interferon response. In addition to the acute inflammatory response, these mice develop rapid SARS-CoV-2 S-specific IgG and neutralizing antibodies between 4- and 7-DPI. While none of the mice exhibited infection-related mortality, this may reflect the immune status of the challenged mice, which were 6-12 weeks old healthy adult mice. Future studies in aged, immunocompromised, and/or obese mice may elicit more dramatic phenotypes, similar to the effect of these comorbidities in human COVID-19 disease.

To better understand the role of type I interferon signaling we infected both IFNAR^−/−^ and IRF3/7^−/−^ and showed that despite similar kinetics of viral clearance, there was significant blunting of ISG response, suggesting that viral replication is resistant to interferon signaling as was previously show with SARS-CoV-1^14^ and has been suggested in SARS-CoV-2 given its vast array of interferon antagonists^19^. While endogenous type I interferon signaling seems to have little effect on SARS-CoV-2 replication, it clearly orchestrates a proinflammatory immune response and may cause immunopathology in patients. Type I interferon signaling is important in protecting against viral infections^20^. Despite the use of type I interferon in the treatment of both MERS and SARS-CoV-1, its endogenous role is less clear with reports of protective effects for MERS^20^ and pathogenic effects for SARS-CoV-1^14^. Notably, overactive or unregulated interferon signaling causes pathology in influenza^21,22^, congenital Zika^23^, and SARS-CoV-1^14,24^ infection. The role of type I interferon signaling in SARS-CoV-2 and COVID-19 is critically important to understanding the protection provided by both the innate and adaptive arms of the immune system and to elucidate any pathogenic effects. These data are especially concerning given current use of type I interferon as a therapeutic for the treatment of COVID-19. Our results indicate a pathological role of type I IFN in COVID-19 respiratory inflammation.

Here we present clear evidence of both the lack of antiviral activity as well as the potential harms of endogenous type I interferon responses in COVID-19 patients. While a similar technique was previously used to develop the first MERS mouse infection system^25^, to our knowledge this is the first report that vector based hACE2 delivery to the mouse respiratory tract is sufficient to enable robust SARS-CoV-2 infection. During the preparation of this manuscript, two publications^5,26^ reported SARS-CoV-2 infection and pathogenesis in different transgenic hACE2 expressing mice. Additionally, infectious models in both hamsters^27^, ferrets^11^ and non-human primates^28^ have also been reported. These models are valuable and have provided much-needed tools for the study of SARS-CoV-2 disease. The hACE2-AAV mouse model described here offers a broadly-available, highly adaptable animal model to understand critical aspects of SARS-CoV-2 viral infection, replication, pathogenesis, with authentic patient derived virus. This model provides a vital platform for testing prophylactic and therapeutic strategies to combat COVID-19.

## Methods

All procedures were performed in a BSL-3 facility (for SARS-CoV-2 infected mice) with approval from the Yale Environmental Health and Safety office.

### Mice

Six to twelve-week-old mixed sex C57Bl/6 (B6J) and B6(Cg)-*Ifnar1*^tm1.2Ees^/J (*Ifnar1*^−/−^) purchased from Jackson laboratories, and IRF3/7 −/− (generous gift from Dr. T. Taniguchi), and were subsequently bred and housed at Yale University. All procedures used in this study (sex-matched, age-matched) complied with federal guidelines and the institutional policies of the Yale School of Medicine Animal Care and Use Committee.

### Antibodies

Anti-I-A/I-E (M5/114.15.2, Pacific blue, B211129, 107620), Anti-CD45 (30-F11, BV605, B283182, 103155), Anti-CD11c (N418, BV711, B265348, 117349), Anti-CD45R/B220 (RA3-6B2, FITC, B230445, 103206), Anti-Ly-6C (HK1.4, PerCP/Cy5.5, B250461, 128011), Anti-CD64 (X54-5/7.1, PE, B270364, 139304), Anti-Ly6G (1A8, PE/Cy7, B194432, 127618), Anti-Siglec-F (E50-2440, A647, 3214986, 562680), Anti-CD11b (M1/70, A700, B259438, 101222), Anti-CD24 (M1/69, APCFire750, B292373, 101840), Anti-CD45.2 (104, Pacific blue, B279653, 109820), Anti-CD4 (RM4-5, BV605, B284681, 100548), Anti-CD8 (53-6.7, BV711, B293230,100759), Anti-CD44 (IM7, FITC, B278352, 103006), Anti-CD62L (MEL-14, PERCP5.5, B272105, 104432), Anti-NK1.1 (PK136, PE, B221273, 108708), Anti-CD69 (H1.2F3, PE/Cy7, B253212, 104512), Anti-CD3 (17A2, APC/Cyanine7, B283940, 100222) antibodies were purchased from BioLegend (Bxxxxxx) or BD Biosciences (xxxxxxx).

### AAV infection

Adeno-associated virus 9 encoding hACE2 were special ordered and purchased from Vector biolabs (AAV-hACE2). Animals were anaesthetized using a mixture of ketamine (50 mg kg^−1^) and xylazine (5 mg kg^−1^), injected intraperitoneally. The rostral neck was shaved and disinfected. A 5mm incision was made and the salivary glands were retracted, and trachea was visualized. Using a 32g insulin syringe a 50μL bolus injection of 10^11^GC of AAV-CMV-hACE2 or control (AAV-GFP or PBS) was injected into the trachea. The incision was closed with 4-0 Vicryl suture. Following intramuscular administration of analgesic (Meloxicam and buprenorphine, 1 mg kg^−1^), animals were placed in a heated cage until full recovery.

### Generation of SARS-CoV-2 Stocks

To generate SARS-CoV-2 viral stocks, Huh7.5 cells were inoculated with SARS-CoV-2 isolate USA-WA1/2020 (BEI Resources #NR-52281) to generate a P1 stock. To generate a working VeroE6, cells were infected at an MOI 0.01 for four days to generate a working stock. Supernatant was clarified by centrifugation (450g x 5min) and filtered through a 0.45 micron filter. To concentrate virus, one volume of cold (4 °C) 4x PEG-it Virus Precipitation Solution (40% (w/v) PEG-8000 and 1.2M NaCl) was added to three volumes of virus-containing supernatant. The solution was mixed by inverting the tubes several times and then incubated at 4 °C overnight. The precipitated virus was harvested by centrifugation at 1,500 x g for 60 minutes at 4 °C. The pelleted virus was then resuspended in PBS then aliquoted for storage at −80°C. Virus titer was determined by plaque assay using Vero E6 cells.

### SARS-CoV-2 infection

Mice were anesthetized using 30% v/v Isoflurane diluted in propylene glycol. Using a pipette, 50μL of SARS-CoV-2 (3×10^7^ PFU/ml) was delivered intranasally.

### Viral RNA analysis

At indicated time points mice were euthanized in 100% Isoflurane. ~33% of total lung was placed in a bead homogenizer tube with 1ml of PBS+2%FBS. After homogenization 250ul of this mixture was placed in 750ul Trizol LS (Invitrogen), and RNA was extracted with RNeasy mini kit (Qiagen) per manufacturer protocol. To quantify SARS-CoV-2 RNA levels, we used the Luna Universal Probe Onestep RT-qPCR kit (New England Biolabs) with 1 ug of RNA, using the US CDC real-time RT-PCR primer/probe sets for 2019-nCoV_N1.

### Viral titer

Lung homogenates were cleared of debris by centrifugation (3900g for 10 minutes). Infectious titers of SARS-CoV-2 were determined by plaque assay in Vero E6 cells in MEM supplemented NaHCO_3_, 4% FBS 0.6% Avicel RC-581. Plaques were resolved at 48hrs post infection by fixing in 10% formaldehyde for 1 hour followed by staining for 1 hour in 0.5% crystal violet in 20% ethanol. Plates were rinsed in water to visualize plaques.

### PRNT assay

Serum samples were heat-inactivated by incubation at 56 ℃ for 30 min before use. Mouse plasma was serially 4-fold diluted from 1:4 to 1:4096, and then an equal volume of SARS-CoV-2 virus was added ~30PFU) and incubated at 37 ℃ for 30minutes. After incubation, 100 ul mixtures were inoculated onto monolayer Vero E6 cells in a 12-well plate for 1 hour. Cells were overlayed with MEM supplemented NaHCO_3_, 4% FBS 0.6% Avicel mixture. Plaques were resolved at 48hrs post infection by fixing in 10% formaldehyde for 1 hour followed by staining for 1 hour in 0.5% crystal violet.

### Immunofluorescence microscopy

Tissue was collected and fixed in 4% PFA overnight. Samples were then dehydrated in a 30% sucrose solution. OCT-embedded 10-mm cryostat sections were blocked in 0.1 M Tris-HCl buffer with 0.3% Triton and 1% FBS before staining. Slides were stained for EpCAM (G8.8, Biolegend) with fluorochrome-labeled primary antibody. The primary antibodies rabbit anti-SARS-CoV-2 nucleocapsid (GeneTex) and rabbit anti-ACE2 (Abcam) or rabbit IgG isotype control were used and detected with secondary antibodies donkey anti-rabbit IgG Alexa Fluor Plus 488/555 (Invitrogen). Slides were stained with DAPI (Sigma) and mounted with Prolong Gold Antifade reagent (Thermo fisher). All slides were analyzed by fluorescence microscopy (BX51; Olympus) with 10x lens. Imaging data were analyzed with Imaris 7.2 (Bitplane).

### Immunohistochemistry

Yale pathology kindly provided assistance with embedding, sectioning and H&E staining of lung tissue. A pulmonary pathologist reviewed the slides blinded and identified immune cell infiltration and other related pathologies.

### Enzyme-linked immunosorbent assay

ELISAs were performed as previously described^9^. In short, Triton X-100 and RNase A were added to serum samples at final concentrations of 0.5% and 0.5mg/ml respectively and incubated at room temperature (RT) for 3 hours before use to reduce risk from any potential virus in serum. 96-well MaxiSorp plates (Thermo Scientific #442404) were coated with 50 μl/well of recombinant SARS Cov-2 S1 protein (ACROBiosystems #S1N-C52H3-100ug) at a concentration of 2 μg/ml in PBS and were incubated overnight at 4 °C. The coating buffer was removed, and plates were incubated for 1h at RT with 200μl of blocking solution (PBS with 0.1% Tween-20, 3% milk powder). Serum was diluted 1:50 in dilution solution (PBS with 0.1% Tween-20, 1% milk powder) and 100μl of diluted serum was added for two hours at RT. Plates were washed three times with PBS-T (PBS with 0.1% Tween-20) and 50μl of mouse IgG-specific secondary antibody (BioLegend #405306, 1:10,000) diluted in dilution solution added to each well. After 1h of incubation at RT, plates were washed three times with PBS-T. Plates were developed with 100μl of TMB Substrate Reagent Set (BD Biosciences #555214) and the reaction was stopped after 15 min by the addition of 2 N sulfuric acid. Plates were then read at a wavelength of 450 nm and 570nm.

### Isolation of mononuclear cells and flow cytometry

Tissue was harvested and incubated in a digestion cocktail containing 1 mg ml−1 collagenase A (Roche) and 30 μg ml−1 DNase I (Sigma-Aldrich) in RPMI at 37 °C for 45 min. Tissue was then filtered through a 70 μm filter. Cells were treated with ACK buffer, and resuspended in PBS with 1% BSA. At this point cells were counted using an automated cell counter (Thermo fisher). Mononuclear cells were incubated on ice with Fc block and Aqua cell viability dye for 20 minutes. After washing, primary antibody staining was performed on ice for thirty minutes. After washing with PBS, cells were fixed using 4% PFA. Cell population data were acquired on an Attune NxT Flow Cytometer and analyzed using FlowJo. Software (10.5.3, Tree Star)

### RNA-seq

Libraries were made with the kind help of Yale Center for Genomic Analysis (YCGA). Briefly, libraries were prepared with an Illumina rRNA depletion kit and sequenced on a NovaSeq. RNA-seq data was aligned using STAR (STAR/2.5.3a-foss-2016b, mm10 assembly) with parameters: --runThreadN 20 --outSAMtype BAM SortedByCoordinate --limitBAMsortRAM 35129075129 --outFilterMultimapNmax 1 --outFilterMismatchNmax 999 -- outFilterMismatchNoverLmax 0.02 --alignIntronMin 20 --alignIntronMax 1000000 -- alignMatesGapMax 1000000 for mapping of repetitive elements. Counts were counted using BEDTools (BEDTools/2.27.1-foss-2016b), coverageBed function, normalized using DESEQ2 and graphed using broad institute Morpheus web tool. For interferon stimulated gene identification, www.interferome.org was used with parameters -In Vivo, -Mus musculus, -fold change up 2 and down 2.

### Graphical Illustrations

Graphical illustrations were made with Biorender.com

## Data availability

RNA-Seq data will be available in SRA upon publication.

## Acknowledgements

This study was supported by awards from National Institute of Health grants, 2T32AI007517-16 (to BI), T32GM007205 and F30CA239444 (to ES), AI054359 and AI127429 (to AI), T32AI007019 (to TM),K08 AI128043 (to CBW), as well as Women’s Health Research at Yale Pilot Project Program (AI, AR), Fast Grant from Emergent Ventures at the Mercatus Center (AI, ES), Mathers Foundation (AR, CBW, AI), and the Ludwig Family Foundation (AI, AR, CBW). A.I. is an investigator of the Howard Hughes Medical Institute.

## Author Contributions

B.I., E. S. and A.I. planned the project. B.I., E.S., T.M. and A.I. designed, analyzed and interpreted data. B.I. and A.I. wrote the manuscript. B.I., E.S., T.M., P.L, A.M and F.L, performed experiments and analyzed data. H.D. bred and cared for animals. R.H., A.R. and C.B.W. provided expertise, materials and analysis of data.

## Competing Interests

None of the authors declare interests related to the manuscript.

## Materials & Correspondence

Correspondence and material requests should be addressed to

## Notes

### Competing Interest Statement

The authors have declared no competing interest.

