## Supplemental figures for "Mouse model of SARS-CoV-2 reveals inflammatory role of type I interferon signaling"

### Extended Data s1-s3

**Extended Data Fig 1. AAV-hACE2 infection makes WT mice susceptible to SARS-CoV-2.** **a,** Immunofluorescence staining of ACE2 in mice infected with AAV-hACE2, 20 days post infection. **b**, Flow cytometry gating strategy for figures **1g**, **3g-i**, extended data fig **1c**, and extended data fig **3a-c**. **c,** Representative flow cytometry plots for figure 1g T cells and NK cells. **d**, Representative flow cytometry plots for figure **1g** myeloid cells. **e**, Different macrophage populations in the lungs of SARS-CoV-2 infected mice two days post infection by flow cytometry. **f**, Serum of mice were collected 7 days after infection with SARS-CoV-2 and limiting dilutions were made to measure reactivity against S1 protein of SARS-CoV-2 using ELISA. **g**, Serum of mice were collected 14 days after infection with SARS-CoV-2 and limiting dilutions were made to measure reactivity against S1 protein of SARS-CoV-2 using ELISA.

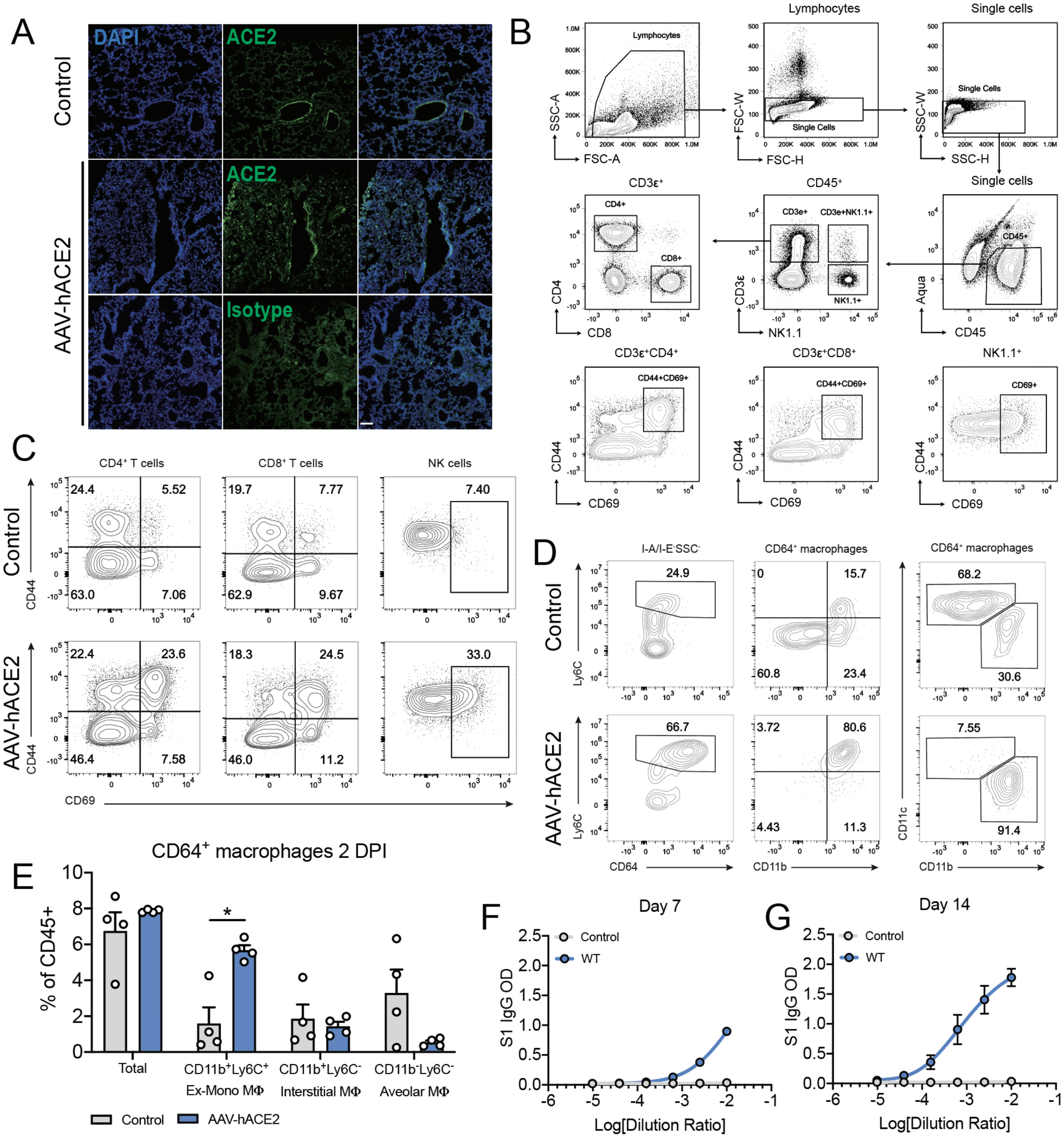

**Extended Data Fig 2. Myeloid cell infiltration in SARS-CoV-2 infected mice. a**, Flow cytometry gating strategy for figures **1g**, **3d-f**, and extended data fig **1b, 1e** and extended data fig. **2b-d. b,** Relative percentage of different cell populations in different knockout mice infected with SARS-CoV-2 two days post infection. **c**, Representative flow cytometry plots for figure **3d** and **e**. **d**, Relative percentage of myeloid cell populations in wildtype mice infected with AAV-hACE2 at days 2 and 4 post SARS-CoV-2 infection.

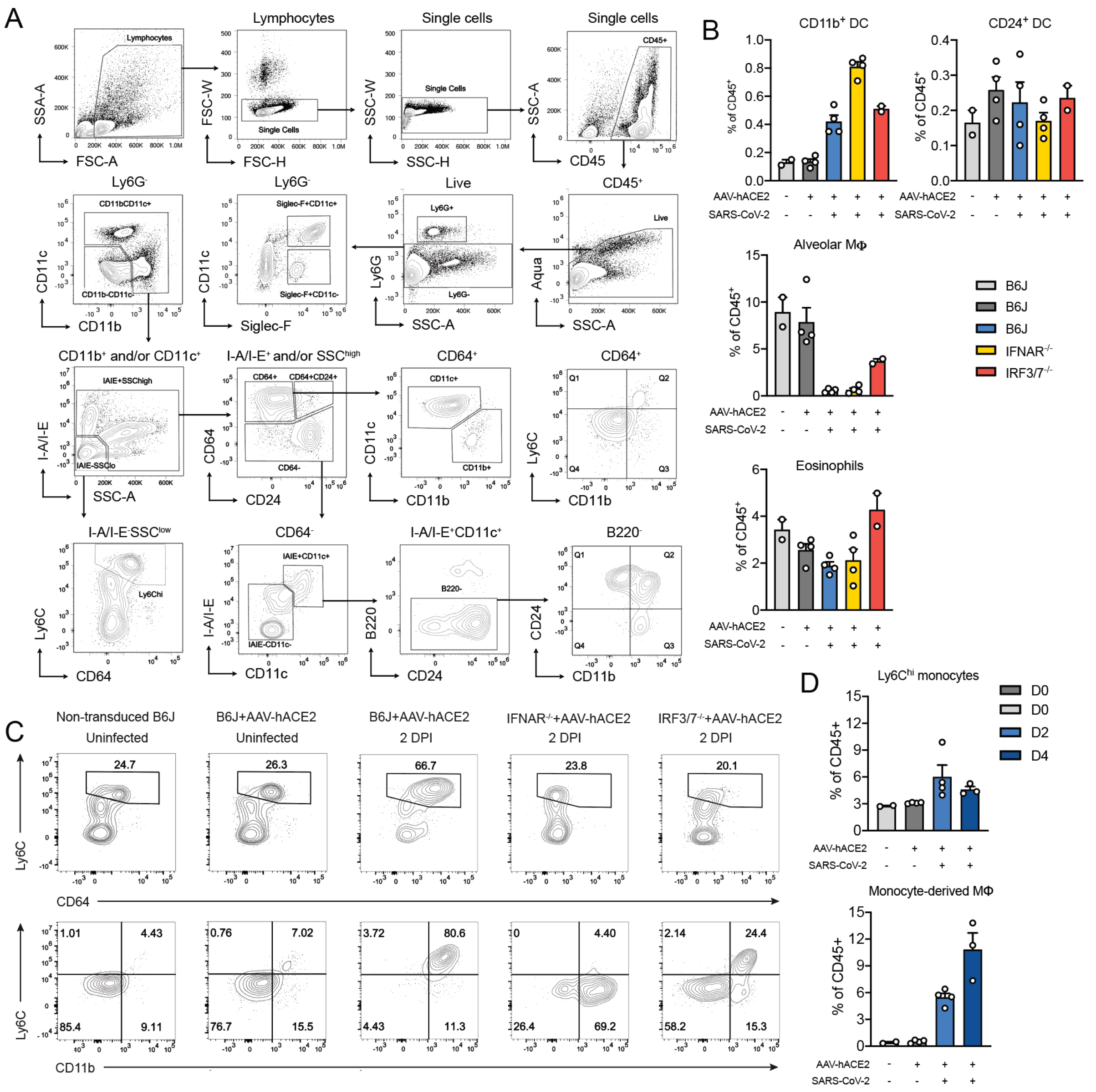

**Extended Data Fig 3. T cell and NK cell infiltration in SARS-CoV-2 infected mice. a**, Representative flow cytometry plots for figure **3g-i**. **b,** Relative percentage of different lymphoid cell populations in different knockout mice infected with SARS-CoV-2 two days post infection. **c**, Relative percentage of lymphoid cell populations in wildtype mice infected with AAV-hACE2 at days 2 and 4 post SARS-CoV-2 infection.

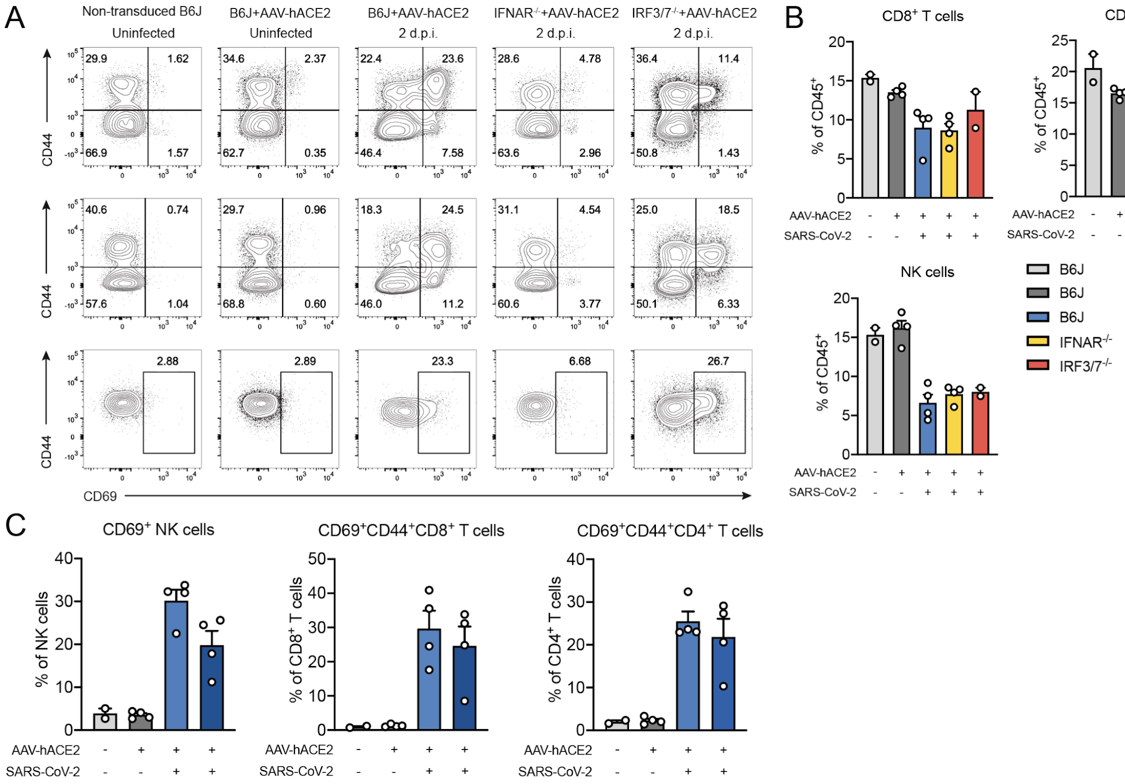
